# BioSAILs: versatile workflow management for high-throughput data analysis

**DOI:** 10.1101/509455

**Authors:** Jillian Rowe, Nizar Drou, Ayman Youssef, Kristin C. Gunsalus

## Abstract

**Motivation:** High-throughput analysis in the current era of systems biology encompasses a range of analysis workflows for different applications, such as transcriptomics, epigenetics, variant discovery, *de novo* genome assembly, etc. Many research institutes house a genomics core facility that is responsible for generating high-throughput sequencing data and often includes a core bioinformatics team that carries out data analysis. Core teams must both keep track of data and maintain numerous software packages and databases – which must be scalable according to ever-changing research needs – and they must allocate computational resources to them. Typical data analysis pipelines involve multiple software packages and require the ability to install, execute, and maintain software stacks on a variety of hardware/operating system configurations, including high-performance computing (HPC) facilities, stand-alone servers, and Cloud services. At the same time, individual researchers need to analyze their data and share their analysis steps with collaborators. Having to rely on *ad hoc* scripts and rigid internally developed pipelines is inefficient, difficult to track and maintain, and ultimately, limits the ability to adapt and evolve as research methods progress.

**Results:** Here, we present BioSAILs (**B**ioinformatics **S**tandardized **A**nalysis **I**nformation **L**ayers), a scientific workflow management system (WMS) developed by the Core Bioinformatics team at NYU Abu Dhabi. BioSAILs comprises two central components, *BioX command* and *HPCRunner command,* supported by *BioStacks* software stacks. BioX structures executable workflows that may be either run directly on a desktop or lab server, or submitted by HPCRunner to HPC infrastructure or the Cloud. BioSAILs is supported by a range of pre-configured, customizable BioStacks software stacks for various applications (e.g. RNA-seq, *de novo* genome/transcriptome assembly, variant discovery, etc.) Software stacks are built using Conda and BioConda (6), have been pre-packaged into modules using EasyBuild, and can be deployed with or without BioSAILs. Within the BioSAILs WMS, users can also invoke commands that take advantage of containerized software in Docker (10) and Singularity (8). Finally, the BioSAILs web resource provides documentation, blog posts related to various BioSAILs analysis tasks, forums, knowledge base & FAQs, as well as an interactive web-based workflow editor/creator. BioSAILs is production-level software that has been in use as the main WMS at NYU Abu Dhabi for the past 2 years and is continuously developed and maintained by the core bioinformatics team.

**Availability:** BioSAILs is open source software, available under the GNU license. Online documentation, support and guides can be found at biosails.abudhabi.nyu.edu. The public Github project is available at Github – BioSAILs. HPC-Runner and BioX-Workflow can be installed through Conda using the BioConda channel.

## BioSAILs Ecosystem

BioSAILs is designed around the idea that a Scientific Workflow Management System should not be a monolithic black box, but should instead incorporate a service-based architecture approach, built with Application Programming Interfaces (APIs). This approach is especially relevant in the quickly changing world of scientific analysis and sequencing technologies, where one size will never fit all. BioSAILs includes APIs to interact with databases (SQLite, MySQL, PostgreSQL), logging services (ElasticSearch), project management tools (JIRA, Redmine), version control systems (Git, GitLab), custom REST APIs, and many others. It aims to be accessible to a wide spectrum of users, from researchers with little to no computational experience, to software engineers looking to extend core functionality.

The BioSAILs framework (Figure 1; see Supplementary Information, BioSAILs Design) comprises central Core components *(BioX* and *HPCRunner* commands) and auxilliary Support components (YAML *Templates, BioStacks* software stacks). The BioX templates contains instructions for analysis workflows. These instructions are interpreted by BioX, which creates a model of the analysis workflow as an executable shell script. HPCRunner then executes the rules specified in the shell scripts as HPC jobs or by submission to a Cloud computing service.

**Figure 1:**
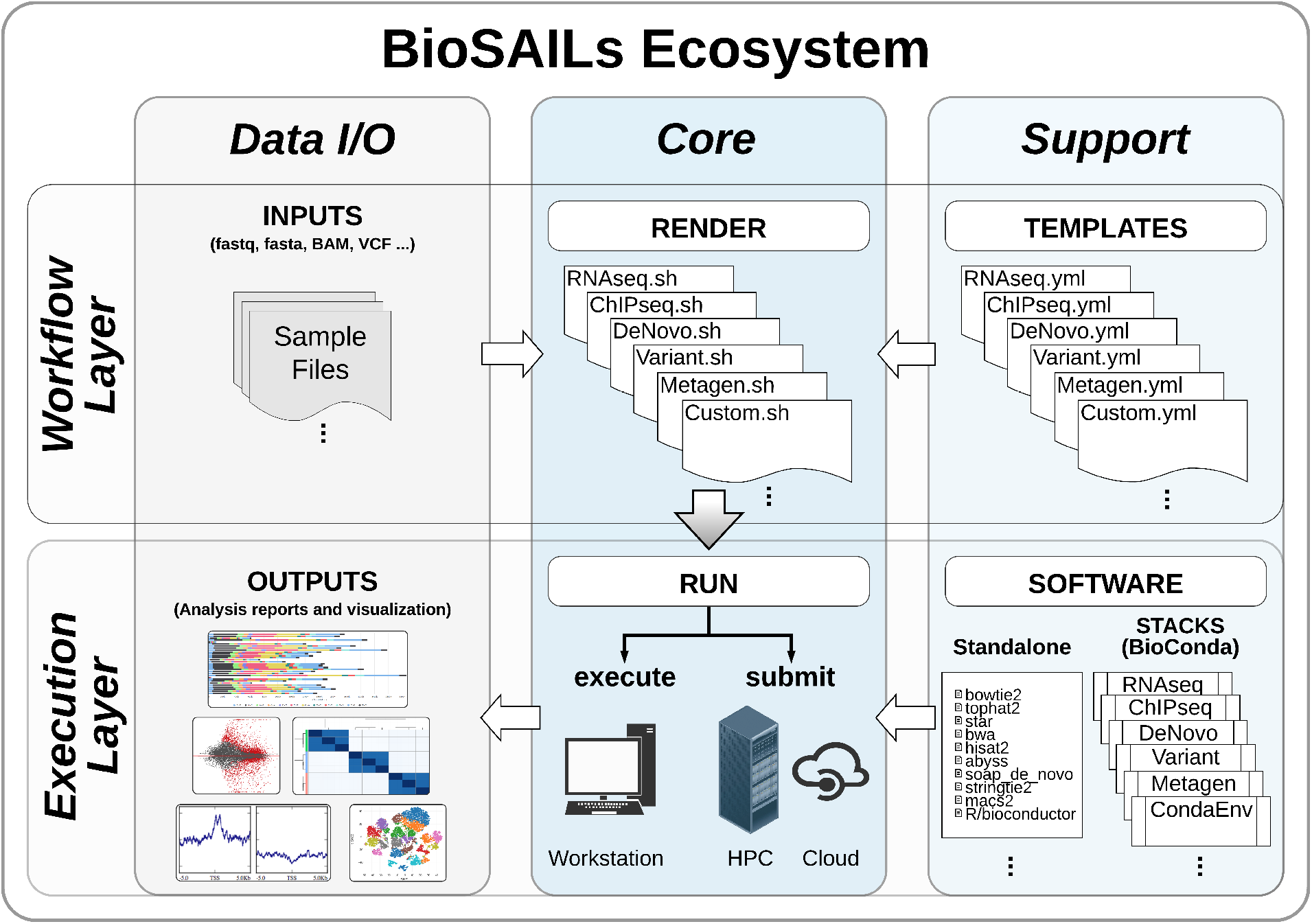
BioSAILs is structured to provide essential Core functions and modular, customizable Support functions. The workflow *TEMPLATES* contains all variables and analysis tasks in YAML format. Templates are *RENDER*ed by the *BioX* command into executable bash scripts, which are then *RUN* by one of two methods: either by direct execution on a workstation, or using the *HPCRunner* command to submit them as workflows to a local HPC cluster or to a Cloud computing platform such as Amazon Web Services (AWS). Users have the option to deploy one or more *SOFTWARE* resources using either *BioStacks* or standalone software.

### Key Features

Key features of BioSAILs include extensibility through plugins, extensive web resources, logging, reporting, and history, and reproducibility through use of bash and caching runs and rendered workflows.

Due to its flexibility, and the fact that BioSAILs has been engineered with the concept of supporting bioinformatics teams, many other functionalities have been built around it, all of which are documented in detail at biosails.abudhabi.nyu.edu. Below, we highlight some key functionalities and “plugins” that BioSAILs supports.

### BioStacks

Software Analysis stacks are built and deployed through a custom layer on top of the Conda (4) package manager. The deployment system is heavily inspired by the BioConda build system. Each BioStack is tailored to a specific use case, either an analysis type (rnaseq, variant detection, de novo genome etc.) or an analysis project. Each stack is under continuous integration and tested on CircleCI (3). Each in-house developed workflow uses one or more of these software stacks, but also allows for users to incorporate their own software, either with Conda or other methods.

### Reproducibility, logging and reporting

The largest gain to reproducible research is having a clear and concise configuration template. BioX moves all the logic of finding files and organizing the workflow structure to the BioX libraries, giving the end user and any collaborators a format that is easy to read and parse. Tasks are implemented as shell (bash) commands, as opposed to a DSL (Domain Specific Language) or programming language. This design choice ensures transparency and maximum support across multiple bioinformatics teams and HPC platforms. Support staff do not need to have any knowledge of BioSAILs in order to help a researcher debug a workflow.

Although users can specify their input/output structure, BioX has a set of best practices for organizing data and directories, making data analysis structure uniform across projects. This allows for an easily understood convention for scientists and collaborators and across project types. These can be viewed in practice through our production workflows, or on the website. Logging is done per submission, per job, and per task with both a tabular terminal view and a programmatic JSON output. Exit codes, duration, stdout, stderr, and original commands are all logged.

### Web Resources and Support

BioSAILs is well documented with a wealth of support and information available online at the BioSAILs website, biosails.abudhabi.nyu.edu. Technical documentation of all the different components of BioSAILs can be found in the Documentation section of the website. More advanced learning topics on how to use BioSAILs for different types of applications can be found in the Blog section. For troubleshooting and other support information, users can check the Knowledge Base section and also post questions on our community-based Forums. The website includes a section on Workflows that contains various types of our in-house developed and community-developed workflows, and additionally hosts an online tool for building, viewing, and editing workflows and then downloading the workflow to a local computer/HPC. The website also contains specific sections dedicated to additional bioinformatics infrastructure resources and how to integrate them with BioSAILs, such as our BioStacks software stacks.

### Plugins

BioSAILs makes heavy use of plugins for customizing functionality and is built to be very developer-friendly. Any function in either BioX or HPCRunner can be overwritten, or modified, using before and after hooks. Important functions are documented in the developer documentation. BioSAILs is written in Perl but is meant to be extensible to developers of other languages. It creates and consumes YAML and JSON files, which are parsable by any programming language (see Supplementary Information, Extending the API for an example extension).

## Comparison with Other Workflow Management Systems

BioSAILs has many similarities to other workflow management systems, and key differences that are outlined in its design philosophy. A full comparison of existing WMSs is beyond the scope of this article and has been addressed previously (9); here we highlight a few of the key differences that distinguish BioSAILs from SnakeMake (7), BcBio (2), BigDataScript (5), and NextFlow (12) (see Table 1).

**Table 1:** WMS Software Comparison

|  | BioSAILS | SnakeMake | BcBio | NextFlow | BigDataScript |
| --- | --- | --- | --- | --- | --- |
| **Usage** |  |  |  |  |  |
| Requires knowledge of DSL to use | No <sup>2</sup> | No <sup>2</sup> | No<sup>2</sup> | Yes<sup>1</sup> | Yes<sup>1</sup> |
| Requires knowledge of DSL to customize <sup>7</sup> | No <sup>3</sup> | No <sup>3</sup> | No<sup>4</sup> | Yes | Yes |
| Requires knowledge of scripting language to customize | No <sup>3</sup> | No <sup>3</sup> | Somewhat <sup>4</sup> | Yes | Yes |
| HPC Schedulers <sup>9</sup> | Yes | Yes | Yes | Yes | Yes |
| AWS Integration | Yes | Yes | Yes | Yes | Yes |
| Google Cloud Integration | Pending | Yes | Yes | Yes | Yes |
| Explicit git / version control integration | Yes | No <sup>5</sup> | No <sup>5</sup> | Yes | No <sup>5</sup> |
| Built in software deployment | Yes <sup>6</sup> | Yes | Yes | No | No |
| **Resources** |  |  |  |  |  |
| Web GUI Workflow Creation | Yes | No <sup>8</sup> | No | No | No |
| Documentation / Web Resources | Yes | Yes | Yes | Yes | Yes |

- Both Nextflow and BigDataScript explicitly state that they are programming languages - BioSAILS, SnakeMake, and BcBio each use templates for workflow creation. A user must be generally familiar with what is a variable and what is text within a template, but not how the template is implemented. - Some knowledge of Perl (BioSAILS) and Python (Snakemake) is helpful, especially concepts such as for loops, but a user can achieve some level customization without learning perl or python. - BcBio comes as a ready made analysis solution. Additional logic must be coded in bcio modules in python. - While any of the scripts and configurations can be kept in a git project root, none but BioSAILS and Nextflow explicitly integrate with github to allow for history and versioning, but a user can implement this on their own in any framework with careful git commits. - In house production workflows have a software stack publically available as a conda env and as a docker container. There is extensive documentation for creating your own through the BioSAILS site and conda documentation. Snakemake and BcBio each provide wrappers around the conda package manager, which BioSAILS does not. - Customization is a very broad term. For more complex operations such as nested looping one needs some knowledge of perl (BioX) or python (Snakemake) - There is a desktop GUI for Snakemake, but it is not implemented by the core snakemake group. - HPC Schedulers include SLURM, SGE, and PBS <sup>6</sup>

Among the different WMSs, BioSAILs is more similar to SnakeMake and BcBio, other template-based WMSs. NextFlow and BigDataScript are each Domain Specific Languages (DSL), and have an audience of software developers or scientists familiar with scripting. Each has community support and extensive documentation. We think BioSAILs has something different to offer with its extensive use of plugins to extend functionality, a workflow creator, github integration, web resources, and extensively researched production workflows for RNA-Seq, Variant Detection, *De Novo* Genome and Transcriptome Assembly, etc.

## Conclusion

Here, we have described BioSAILs, an open source scientific workflow management system capable of supporting multiple analysis tasks that can integrate seamlessly with existing computational infrastructures, Cloud services, or standalone workstations. It is transparent, highly customizable, includes a host of API support, and can be easily extended. Copious logging and reporting means that BioSAILs has been implemented with analysis best practices in mind, capable of delivering reproducible research. The software benefits from web resources that provide extensive documentation and support through the BioSAILs website, making it easier for users to adopt.

## Acknowledgments

The authors would like to acknowledge the Easybuild and Conda/BioConda teams for the excellent work on software packaging and deployment and the NYU Abu Dhabi High Performance Computing department for providing the necessary resources for the development of BioSAILs. We would also like to acknowledge the Advanced Computing WCMCQ, where the first releases of the biox and hpcrunner commands were initially implemented. This work was supported by a grant from the NYU Abu Dhabi Research Institute to the NYU Abu Dhabi Center for Genomics & Systems Biology (CGSB). Finally, the authors would like to acknowledge all the faculty and researchers at the NYU Abu Dhabi CGSB and Biology division for their excellent work, which has motivated the development of BioSAILs.

