## Supplementary for "BioSAILs: versatile workflow management for high-throughput data analysis"

### Supplementary Information

#### BioSAILs Design

##### BioSAILs workflow template

The configuration template is the set of instructions that the users want to execute. In essence, it provides all the necessary parameters, analysis steps, input/output structures, and definition of computational resources to be used. The templates are written in YAML format and consist of three main sections: samples, variables, and rules. Rules consist of the actual commands that are executed (e.g. samtools, some type of alignment software, quality trimming, etc). The template (detailed in Supplementary Information) is then processed by BioX (see Supplementary Information, BioX Example Workflow) and executed on a workstation, HPC, or cloud with HPCRunner.

##### BioX Command

BioX comprises the set of libraries, along with their commandline interfaces, that convert a workflow template to an executable shell script which can be either run as is, or executed using HPCRunner. BioX has a number of helper functions for finding samples and looping over them, and is designed to provide enough defaults to get a user started. Each aspect of the workflow, from declaring variables to creating process templates, can be customized by including outside configuration files (yaml, json, or ini), by exporting rule data to JSON for use with another language (Python, Ruby, R), or by including custom Perl modules. Each time a BioX workflow is processed a "frozen" copy is kept in the ".biosails" cache directory, which includes the datetime of processing along with any additional commandline parameters. This, in conjunction with HPCRunner's git integration, gives a detailed history of the computation and analysis of a given project. Each time a workflow is executed with HPCRunner a user can optionally add a commit message, and if a project is under git integration a version per execution will be created. History can be browsed visually using any git GUI, or on a terminal using the hpcrunner.pl "history" command. Current BioX plugins include a web-based workflow creator written in html and Angular.js (1) that exposes some functions to a REST API ( BioX-Workflow-Command-REST ).

##### HPCRunner Command

The *biosails* command wraps around both *biox* and *hpcrunner*, and can execute subcommands from either. HPCRunner-Command takes a shell script, along with an optional simple markup, and communicates how that job should be run. This logic is split into three main categories: submitting jobs to a cluster, executing a task, and reporting/logging. For a submission on a cluster, HPCRunner takes the jobs, and their dependencies, and creates a directed graph (see Supplementary Figure 3), which it communicates to the scheduler. This is also done using a template, which the user can customize. Once a job has been submitted, each task within that job is executed in a fashion similar to GnuParallel (11). Each task is executed as a standalone process, but multiple tasks can be executed in parallel by setting the HPC directives. Different combinations of stacking ensure that a researcher can work within the limits of their compute environment. Each task has a corresponding logfile that has important environmental information such as hostname, variables exported by the scheduler such as job and task ID, process ID, start time, completion time, duration, and exit code, along with all standard out and standard error. Files are organized consistently using dates, job names, and task indices, and are uniform across analysis types.

HPCRunner can be extended through the use of additional plugins, which can be passed on the command line through a configuration file. Plugins are used either during the submission, during job execution, or both. For execution, there are plugins to integrate with different databases, mainly SQLite (Logger-Sqlite) and ElasticSearch (Logger-ElasticSearch). Workflows can be exported to JSON for use with any templating language. Currently, there is a Python Jinja2 templating utility included.

Although BioSAILs is highly customizable, in the end, and given an analysis YAML template, a user can execute the analysis simply by initiating two short commands:

```
biosails render -w analysis_template.yml -o analysis_script.sh
biosails submit_jobs --infile analysis_script.sh
```

#### BioSAILs - Technical Architecture

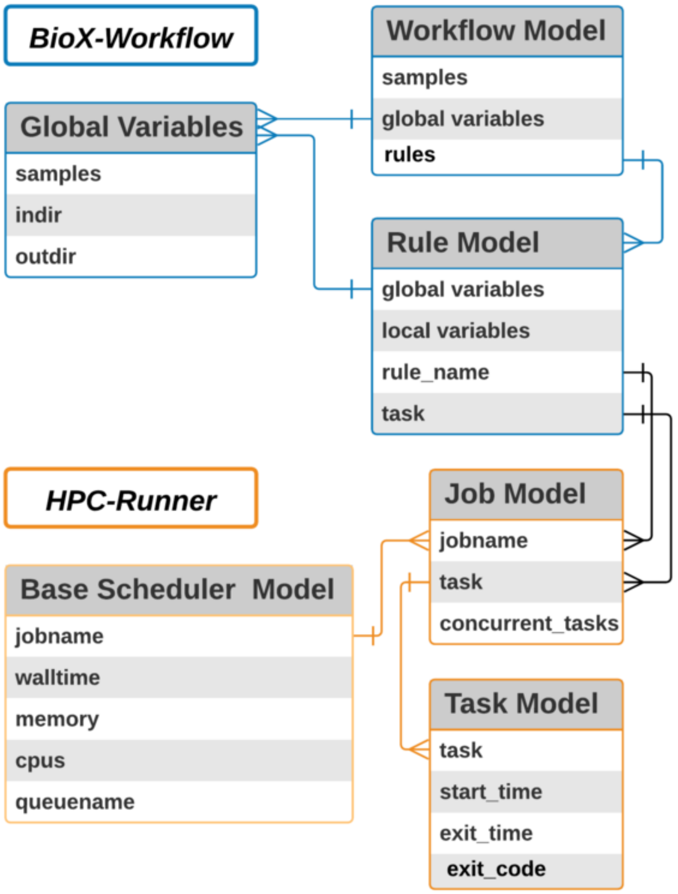


Figure 2: The technical architecture can be described as a set of relationships between the workflow variables (for example reference genome locations, cpu requirements for a job, etc), along with the relationships of the dependencies per job/rule (e.g. alignment depends upon QC) and the individual task.

#### BioX Example Workflow

##### Samples

###### *Example Sample Definition*

```
---
global:
    # Samples Definition
```

```
    ## Find samples in 'indir'
    - indir:   "data/processed"
    ## Find samples in 'indir' matching the 'sample_rule' pattern
    - sample_rule: (Sample.*)$
    ## Samples can be files or directories,
    ## in this case we are stating that each sample is a directory.
    - find_by_dir: 1
    # Put results here
    - outdir:  "data/analysis"
```

***Example Directory Structure***

```
#Example directory Structure
  tree data
data
+ processed
        + Sample_01
                Sample_01_R1.fastq.gz
                Sample_01_R2.fastq.gz
        + Sample_02/
                Sample_02_R1.fastq.gz
                Sample_02_R2.fastq.gz
        + Sample_03/
                Sample_03_R1.fastq.gz
                Sample_03_R2.fastq.gz
```

At the beginning of each workflow are instructions that allow BioX-Workflow to find the samples. The way that BioX deals with samples and integrates them into rules is one of the defining features of BioSAILs, and was a specific design choice for this software. Workflows support the same features YAML supports, including comments and long strings.

##### Variable Declaration

```
---
global:
    #Other variables created here
    ...
    - reference: "/scratch/gencore/references/grch38.fa"
```

Any variables can be declared within either the global or local rules, in any format YAML accepts, or as inline Perl code . Global variables are available anywhere within a workflow, while local variables are only available for that rule. Additional json/yaml files can also be included, but these cannot contain inline Perl code. Additionally, BioX is able to export its data to JSON at any time during the workflow, allowing for integration with other languages and APIs. At present there is a Python integration supported within the library, and a node.js integration supported as an external project. HPC variables are usually those passed to HPCRunner for execution, but are not constrained.

##### Rules Declaration

```
    - bwa_mem:
        local:
                # This is the first rule, so it will inherit the indir from the
                # global indir above (i.e. data/processed/Sample0[123]/), but can be
                # specified locally as well.
```

```
            # -indir: "data/processed/{$sample}/"

            # By default the outdir is also inherited from the global variable
            # plus the rule name. For example for Sample_01, the output directory
            # here will resolve to data/analysis/Sample01/bwa_mem/. Again this can
            # be specified locally to any other path.
            # - outdir: "data/analysis/{$sample}/bwa_mem"
            - HPC:
              - cpus_per_task: 12
              # In this HPC variables section, we only specified the number of
              # CPUs to use, which can be interpreted by an HPC scheduler
              # (SLURM for example). Any other additional computational parameters
              # can also be specified, as long as they can be interpreted by
              # a scheduler, such as memory and walltime etc.
              #- mem: '60GB'
              #- walltime: '12:00:00'
        process: |
            #TASK tags={$sample}
            bwa mem -t 12 -M \
                 {$reference} \
                 {$indir}/{$sample}_R1.fastq.gz \
                 {$indir}/{$sample}_R2.fastq.gz \
                 > {$outdir}/{$sample}_aligned.sam
    - samtools_view:
        local:
            # Indir is inherited from the previous rule (outdir), but can be
            # specified, or overwritten, by the user as a "local variable".
            # In this example, the indir in the samtools_view rule will
            # automatically be resolved as data/analysis/Sample_0[123]/bwa_mem/
            - HPC:
              - deps: 'bwa_mem'
              - mem: '50GB'
              - walltime: '14:00:00'
        process: |
            #TASK tags={$sample}
            samtools view -b -S {$indir}/{$sample}_aligned.sam \
                > {$outdir}/{$sample}_aligned.bam
```

Variables can reference other variables, be scalar (strings or integers), lists, or objects, and/or have embedded Perl code. Curly braces mark the beginning of a variable. BioX-Workflow encourages scientists to use best practices by clearly outlining inputs and outputs, but these are not necessary and could be referenced directly. HPC variables are included directly within a workflow for execution context. At any point the data within a rule can be exported to YAML or JSON, two commonly used configuration file types, for use with other languages (Python, Ruby, Go, etc).

Variable declaration can be as simple or as complex as a user desires. Variables can reference other variables, or be grouped together into particular data structures. A commonly used convention at NYUAD is to group variables into INPUT and OUTPUT (see Supplementary Production Workflow).

##### HPC Directives

```
- bwa_mem:
        local:
```

```
                ...
                - HPC:
                  - modules: 'gencore gencore_biosails gencore_variant_detection'
                  - mem: '60GB'
                  - cpus_per_task: 12
                  - walltime: '12:00:00'
        ...
    - samtools_view:
        local:
                ...
                - HPC:
                  - modules: 'gencore gencore_biosails gencore_variant_detection'
                  - deps: 'bwa_mem'
                  - mem: '50GB'
                  - walltime: '14:00:00'
        ...
```

HPC directives are generalized to an easy to use and remember syntax that remains the same across schedulers. It is then parsed to the format that the scheduler understands. Support for each scheduler (SLURM, PBS, SGE, AWS Batch) is implemented as a plugin and can be customized.

HPC modules can be any module supported by environment modules or lmod. Each of the production workflows produced by NYUAD has one or more accompanying software stacks, BioStacks. These are available publicly as Conda environments through Anaconda Cloud, or as docker images through quay.io. Each BioStack can be deployed as a traditional HPC or as a conda environment. Users can create their own Conda environment and source those either inline with the task, or using the HPC directive 'conda_env'. Additionally, any software available in a user's shell is available to BioSAILs with no special configuration.

##### Process Declaration

The process is the final product of a rule. It is rendered to bash as the task that is executed by HPC-Runner.

```
        process:  |
                #TASK tags={$sample}
                bwa mem -t 12 -M \
                     {$reference} \
                     {$INPUT}_read1_trimmomatic_1PE.fastq.gz \
                     {$INPUT}_read2_trimmomatic_2PE.fastq.gz \
                     > {$OUTPUT}
```

Each of the variables is evaluated to produce a task. The default behavior is one task per sample, but this behavior can be overridden or customized within the workflow itself, by using plugins, or by creating custom Perl modules.

```
#TASK tags={$sample}
bwa mem -t 12 -M \
    {$reference} \
    Sample_01_read1_trimmomatic_1PE.fastq.gz \
    Sample_01_read2_trimmomatic_2PE.fastq.gz \
        > Sample_01.aligned.sam
```

##### Template Rendering

Once a template is complete, the command "biox" run renders it, and also checks for common mistakes such as not finding samples, undeclared variables, syntax errors, etc.

```
#Run the entire workflow
biosails render -w workflow.yml -o workflow.sh
#Select particular rules from the workflow
biosails render -w workflow.yml -o workflow.sh --select_rules bwa_mem
```

###### Execution

Once a BioX template is complete it can be rendered as a regular bash script. Each workflow is annotated with arguments that were used to create it along with date and time of rendering. The bash script can be run as is, or executed with HPCRunner, or submitted to a cluster using HPCRunner.

```
# Submit to a cluster (Default is Slurm, but other schedulers can be chosen)
biosails submit_jobs --infile workflow.sh
# Execute directly on a workstation or compute node
biosails execute_node --infile workflow.sh
```

#### HPCRunner Command

##### Directed Graph

A series of analysis rule can be computationally described as a directed graph, where each rule in the workflow is dependent upon zero or more rules. For instance, in our production variant calling workflow, we initially split bam files by chromosome after alignment for faster processing, and then combined across samples per chromosome for the joint genotype calling (gvcf) vcf. In the portion of the workflow shown here the first rule is 'combine_gvcfs_per_chromosome' .

#### BioX-Workflow Command

##### Example Workflow

```
---
global:
    # Initial Directory Setup
    - indir:   "data/processed"
    - outdir:  "data/analysis"
    # Find Samples
    - sample_rule: (Sample.*)$
    - find_by_dir: 1
    # Output Directory Structure
    - by_sample_outdir: 1
    # Processed Dirs
    - trimmomatic_dir: "data/processed/{$sample}/trimmomatic"
    - HPC:
       - account: 'gencore'
       - partition: 'serial'
       - module: 'gencore gencore_biosails gencore_variant_detection/1.0'
       - cpus_per_task: 1
       - commands_per_node: 1
rules:
  - bwa_mem:
      local:
        - indir: '{$trimmomatic_dir}'
        - outdir: '{$bwa_mem_dir}'
        - INPUT: '{$indir}/{$sample}'
```

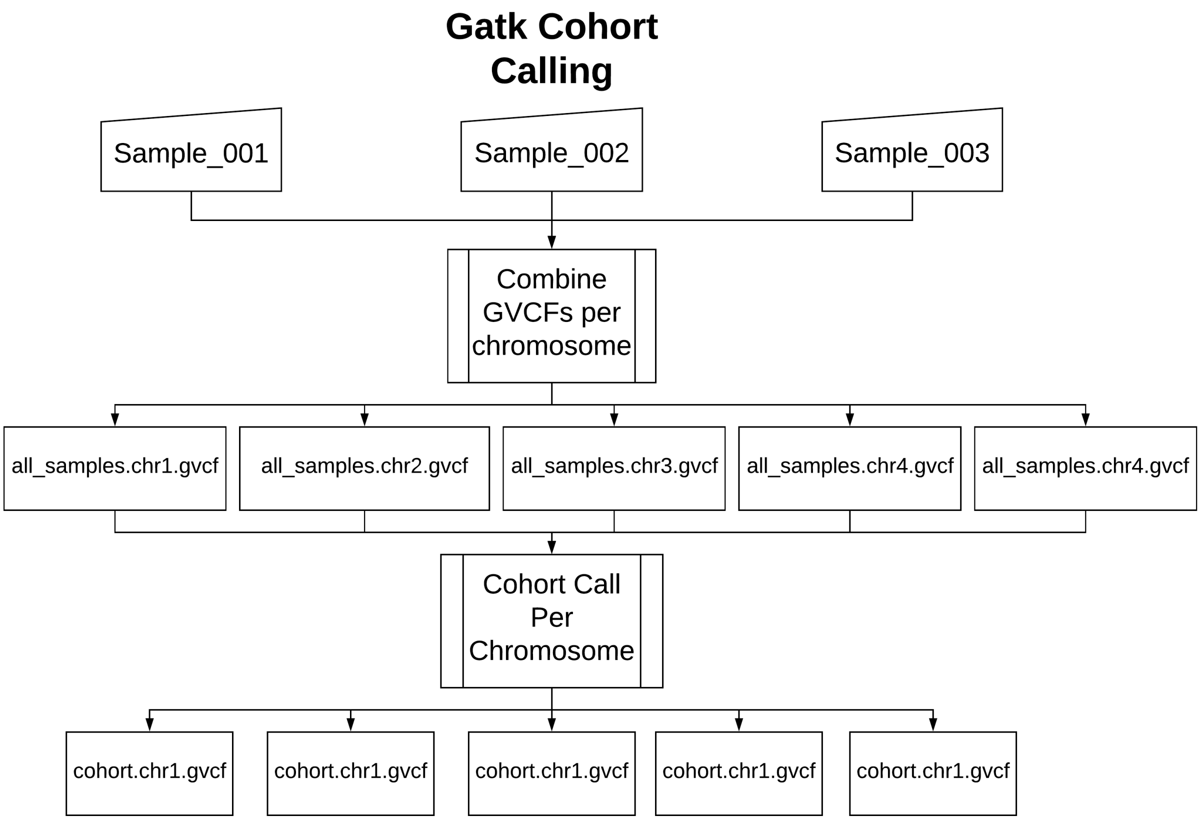


Figure 3: Directed Graph - GATK Cohort Calling

```
    - OUTPUT: '{$outdir}/{$sample}_aligned.sam'
    - HPC:
      - mem: '60GB'
      - cpus_per_task: 12
      - walltime: '12:00:00'
  process: |
    #TASK tags={$sample}
    bwa mem -t 12 -M \
         {$reference} \
         {$INPUT}_read1_trimmomatic_1PE.fastq.gz \
         {$INPUT}_read2_trimmomatic_2PE.fastq.gz \
         > {$OUTPUT}
```

#### HPCRunner Command

##### HPCRunner Stats

Logging plugins are currently implemented for ElasticSearch and SQLite, with plans to extend to MySQL and Postgres. The core HPCRunner libraries implement an interface that is shared across logging interfaces. This makes it possible to extend the features to databases, search services, REST APIs, project management system, etc. Each logger has a terminal output for human readability as well as a JSON output for use with outside services. Internally, BioX/HPCRunner is used with JIRA in order to track workflows. HPCRunner can optionally integrate with git in order to track workflow and submission versions. Status updates can be rendered directly on the command line.

```
> hpcrunner.pl stats --data_dir \
    /home/gencore/hpcrunner-test/.biosails/.hpcrunner-data/job1/
.---------------------------------------------------------.
|        Time: 2017-08-03T18:11:49 Project: job1          |
|   SubmissionID: AB96B038-7855-11E7-9A85-F40564667706    |
+---------+----------+---------+---------+------+-------+
| JobName | Complete | Running | Success | Fail | Total |
+---------+----------+---------+---------+------+-------+
| test1   |        5 |       0 |       5 |    0 |     5 |
| test2   |        5 |       0 |       5 |    0 |     5 |
'---------+----------+---------+---------+------+-------
```

Figure 4: Status Table

```
hpcrunner.pl stats -l --data_dir \
    /home/gencore/hpcrunner-test/.biosails/.hpcrunner-data/job1/
.------------------------------------------------------------------------------------------------------------------------.
|                                        Time: 2017-08-03T18:11:49 Project: job1                                         |
|                                     SubmissionID: AB96B038-7855-11E7-9A85-F40564667706                                 |
+---------+--------+-----------+--------------------+--------------------+-----------------------------------------+----------+
| Job | ID | Task Tags | Start Time          | End Time            | Duration                                | Exit Code |
+---------+--------+-----------+--------------------+--------------------+-----------------------------------------+----------+
| test1   |      1 | s1        | 2017-08-03 18:12:05 | 2017-08-03 18:13:55 |  0 days, 00 hours, 01 minutes, 50 seconds |         0 |
| test1   |      2 | s2        | 2017-08-03 18:12:05 | 2017-08-03 18:13:55 |  0 days, 00 hours, 01 minutes, 50 seconds |         0 |
| test1   |      3 | s3        | 2017-08-03 18:12:05 | 2017-08-03 18:13:55 |  0 days, 00 hours, 01 minutes, 50 seconds |         0 |
| test1   |      4 | s4        | 2017-08-03 18:12:05 | 2017-08-03 18:13:55 |  0 days, 00 hours, 01 minutes, 50 seconds |         0 |
| test1   |      5 | s5        | 2017-08-03 18:12:05 | 2017-08-03 18:13:55 |  0 days, 00 hours, 01 minutes, 50 seconds |         0 |
| test2   |      6 | s1        | 2017-08-03 18:14:18 | 2017-08-03 18:14:28 |  0 days, 00 hours, 00 minutes, 10 seconds |         0 |
| test2   |      7 | s2        | 2017-08-03 18:14:18 | 2017-08-03 18:14:28 |  0 days, 00 hours, 00 minutes, 10 seconds |         0 |
| test2   |      8 | s3        | 2017-08-03 18:14:18 | 2017-08-03 18:14:28 |  0 days, 00 hours, 00 minutes, 10 seconds |         0 |
| test2   |      9 | s4        | 2017-08-03 18:14:19 | 2017-08-03 18:14:29 |  0 days, 00 hours, 00 minutes, 10 seconds |         0 |
| test2   |     10 | s5        | 2017-08-03 18:14:19 | 2017-08-03 18:14:29 |  0 days, 00 hours, 00 minutes, 10 seconds |         0 |
'---------+--------+-----------+--------------------+--------------------+-----------------------------------------+----------
```

Figure 5: Detailed Status Table

##### Variable Declaration

HPCRunner borrows its variable declaration syntax from schedulers such as SLURM and PBS.

```
#HPC jobname=bowtie2
#HPC partition=serial
#HPC mem=24GB
#HPC cpus_per_task=7
#HPC module=gencore_variant_detection

bowtie2 -p 7 -x reference.fa \
-1 sample1.read1.1PE.fastq.gz \
-2 sample1.read2.2PE.fastq.gz -S sample1.sam
```

The '#HPC VAR=VALUE' assigns the value VALUE to VAR. Variables can be added or changed through custom plugins.

#### Task Execution Deep Dive

##### Dependency Declaration

HPCRunner incorporates two levels of directed acyclic graphs, DAGs. The first is between jobs, and the second is within job task dependencies. Task dependencies are declared using 'task tags'.

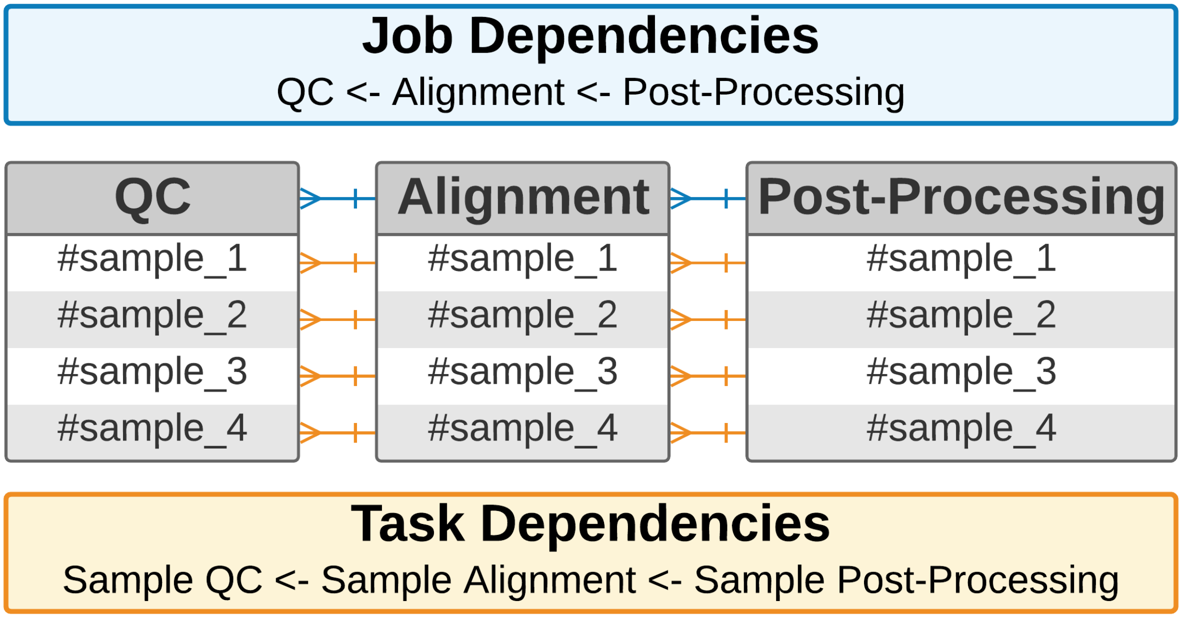


Figure 6: Dependency Declaration Overview

```
#HPC jobname=bowtie2

#TASK tags=sample1
bowtie2 -p 7 -x reference.fa \
-1 sample1.read1.1PE.fastq.gz \
-2 sample1.read2.2PE.fastq.gz \
-S sample1.sam
#TASK tags=sample2
bowtie2 -p 7 -x reference.fa \
-1 sample2.read1.1PE.fastq.gz \
-2 sample2.read2.2PE.fastq.gz \
-S sample2.sam

#HPC jobname=samtools_view
#HPC deps=bowtie2
```

```
#TASK tags=sample1
samtools view -b -S sample1.sam > sample1.bam
#TASK tags=sample2
samtools view -b -S sample2.sam > sample2.bam
```

The 'samtools_view' job has a dependency on the 'bowtie2' job. If there were no task dependencies, all bowtie2 tasks would have to complete before samtools_view tasks would begin, but using task dependencies HPCRunner builds dependencies per task within a job. Each job is submitted as one or more job arrays, and task dependencies are declared using the array ID where possible.

Generally, each task is a separate element in the job array, which we want farmed out as widely as possible, but it can be the case that we want to stack tasks together, particularly with large workflows where we consume more resources than available. In this case we can use the '#HPC commands_per_node=X', which will tell HPCRunner that there are X separate tasks to execute instead of one. HPCRunner can also be told to be its own task manager in a sense, by enabling '#HPC procs=Y' in order to run Y number of tasks in parallel. No matter how many tasks are being executed, each is run in isolation.

#### Extending the API

##### Serving BioX as a REST API for frontend interfaces

On the BioSAILs website we have a front end interface for users to create their own new workflows or to customize existing ones. The server side is a REST API extension on top of BioSAILs. The extension easily wraps around the API, and instead of writing the results out to a bash file as we normally would, transforms the results of the BioSails render call to an object that is then served to the front end interface through the REST API. We left out some error checking in this example to make the code more easily understandable, but the full code is available at BioX-REST.

```
package Main;

use MooseX::App::Command;
#################################################################
## Any of the BioSAILs modules can be extended
#################################################################
extends 'BioX::Workflow::Command::run';
with 'BioX::Workflow::Command::run::Utils::Rules';

option '+workflow' => ( required => 0, );

has 'text_obj' => (
    is      => 'rw',
    isa     => 'HashRef',
    default => sub { {} },
);

#################################################################
## Before, after, and around hook allows developers
## to modify the API at any point
## Here we are modifying plain text string that is returned,
## and putting it into an object, which is returned through the REST API
#################################################################

around 'eval_process' => sub {
```

```
    my $orig = shift;
    my $self = shift;

    my $text = $self->$orig(@_);

    if(! exists $self->text_obj->{$self->rule_name} ){
      $self->text_obj->{$self->rule_name} = [] ;
    }

    push(@{$self->text_obj->{$self->rule_name}}, $text);
};

package BioX::Workflow::Command::REST::run;

use strict;
use warnings;

use Main;
use HTTP::Status qw(:constants);
#################################################################
## Raisin::API is a small perl module that creates a REST API
#################################################################
use Raisin::API;
use Types::Standard qw(HashRef Any Int Str ArrayRef);
use File::Temp qw/ tempfile tempdir /;
use Capture::Tiny ':all';


api_format 'json';
api_default_format 'json';

desc 'Run a Workflow';
#################################################################
## Just as you would pass in command line parameters,
## pass in and describe the REST parameters
#################################################################

resource run => sub {
    summary 'List users';
    params(
        optional(
            'select_rules',
            type => ArrayRef [Str],
            default => [],
            desc    => 'Process one or more rules'
        ),
        optional(
            'samples',
            type => ArrayRef [Str],
            default => [ 'Sample_01', 'Sample_02' ],
            desc    => 'Process one or more samples'
```

```
        ),
        optional(
            'data',
            type => HashRef,
            desc => 'Workflow object',

            #TODO add global and rules here
            group {}
        ),
    );

    post sub {
        my $params = shift;

        my $biox = Main->new();
        my $tmp_dir = tempdir( CLEANUP => 0 );
        chdir($tmp_dir);

        my $samples      = $params->{samples};
        my $data         = $params->{data};
        my $select_rules = $params->{select_rules};

        my ( $stdout, $stderr, $exit ) = capture {
            $biox->select_rules($select_rules);
            $biox->stdout(1);
            $biox->samples($samples);

            $biox->print_opts;
            $biox->workflow_data($data);
            $biox->apply_global_attributes;

            $biox->global_attr->create_outdir(0);
            $biox->global_attr->coerce_abs_dir(0);
            $biox->get_global_keys;

            $biox->write_workflow_meta('start');
            $biox->iterate_rules;
        };

        { process_text => $stdout, logs => $stderr, text_obj => $biox->text_obj };
    };
};

1;
```

[2] BcBio Next Gen.

[3] CircleCI. https://circleci.com.

[4] Conda. https://conda.io/docs/.

[5] P. Cingolani, R. Sladek, and M. Blanchette. BigDataScript: a scripting language for data pipelines. *Bioinformatics*, 31(1):10–16, sep 2014.

[6] Björn Grüning, Ryan Dale, Andreas Sjödin, Jillian Rowe, Brad A. Chapman, Christopher H. Tomkins-Tinch, Renan Valieris, and Johannes Köster and. Bioconda: A sustainable and comprehensive software distribution for the life sciences. oct 2017.

[7] J. Koster and S. Rahmann. Snakemake–a scalable bioinformatics workflow engine. *Bioinformatics*, 28(19):2520–2522, aug 2012.

[8] Gregory M. Kurtzer, Vanessa Sochat, and Michael W. Bauer. Singularity: Scientific containers for mobility of compute. *PLOS ONE*, 12(5):e0177459, may 2017.

[9] Jeremy Leipzig. A review of bioinformatic pipeline frameworks. *Briefings in Bioinformatics*, page bbw020, mar 2016.

[10] Dirk Merkel. Docker: Lightweight linux containers for consistent development and deployment. *Linux J.*, 2014(239), March 2014.

[11] O. Tange. GNU Parallel - The Command-Line Power Tool. Technical report, 2011. Accessed on Wed, September 12, 2018.

[12] Paolo Di Tommaso, Maria Chatzou, Evan W Floden, Pablo Prieto Barja, Emilio Palumbo, and Cedric Notredame. Nextflow enables reproducible computational workflows. *Nature Biotechnology*, 35(4):316–319, apr 2017.
